# Light assisted antibiotics

**DOI:** 10.1101/229914

**Authors:** Aikaterini Argyraki, Merete Markvart, Camilla Stavnsbjerg, Kasper Nørskov Kragh, Lars Bjørndal, Thomas Bjarnsholt, Paul Michael Petersen

**Affiliations:** Department of Photonics Engineering, Technical University of Denmark, Frederiksborgvej 399, DK-4000, Roskilde, Denmark; Cariology and Endodontics, Department of Odontology, Faculty of Health Science, University of Copenhagen, Nørre Allé 20, DK-2200, Copenhagen N, Denmark; Department of Immunology and Microbiology, Costerton biofilm Center, Faculty of Health Science, University of Copenhagen, Blegdamsvej 3B, DK-2200v Copenhagen N, Denmark; Department of Clinical Microbiology, Rigshospitalet, Juliane Maries vej 22, 2100 Copenhagen Ø, Denmark

## Abstract

The overuse of antibiotics is accelerating the bacterial resistance, and therefore there is a need to reduce the amount of antibiotics used for treatment. Here, we demonstrate that specific wavelengths in a narrow range around 296 nm are able to eradicate bacteria in the biofilm state more effectively, than antibiotics and the combination of irradiation and antibiotics is even better, introducing a novel concept *light assisted antibiotics*. The investigated wavelength range was 249 nm to 338 nm with an approximate step of 5 nm. The novel concept can significantly reduce the amount of antibiotics needed for eradicating mature bacterial biofilms. The irradiation treatment was combined with tobramycin and its efficiency was compared to combinatory antibiotic treatment and highly concentrated antibiotic monotherapy. The eradication efficacies, on mature biofilms, achieved by light assisted antibiotic and by the antibiotic monotherapy at 10-fold higher concentration, were equivalent. The present achievement could motivate the development of light assisted antibiotic treatments for treating infections.

---

There is a worldwide increasing awareness and concern about the antibiotic resistance, which in the future could prevent effective treatment of a large number of infectious diseases [1]. The emergency of antibiotic resistance has evoked a “turn” towards antibiotics control/reduction programs [2]. Moreover, biofilm infections are highly persistent to the immune response and known for their tolerance to antibiotic treatments [3]. Alternative approaches, different than conventional antibiotic treatments, are gaining increasing interest and light based treatments [4] are unconventional strategies for biofilm eradication. Bak *et al.* has reported *in vivo* disinfection potential of catheter lumens by UVC LEDs [5]. Recently, we have observed that UVB irradiation is more efficient than UVC, in eradicating biofilms plated on cellulose nitrate membrane filters [6]. However, the ultraviolet (UV) wavelength dependent eradication efficiency for *Pseudomonas aeruginosa* biofilms has not been reported. Apart from UV light, also exposure to blue light has been proven to have antimicrobial effect, without the requirement of exogenous photosensitizers present [7].

UV light emitting diodes (LEDs) are progressing as light source options in biomedical applications mainly due to their flexibility in spectral design and ease in operation. AlGaN LEDs are continuously progressing as UV light sources. The shortest wavelength achieved with pure AlN LEDs is 210 nm [8]. External quantum efficiency (EQE) of UV LEDs is continuously improving as both internal quantum efficiency (IQE) and light extraction efficiency (LEE) are boosted by various techniques like: migration-enhanced metalorganic chemical vapor deposition [9, 10], ammonia pulsed-flow method [11, 12] and nanowires [13], or surface plasmons [14] and aluminum reflective electrodes [15], respectively. EQE above 10% has been reported when approaches for improving IQE and LEE are applied in combination [16].

Combinatory therapies seem to be the solution for combating tolerant biofilms present in chronic infections [17, 18]. Particularly, if the combinatory treatments enact complementary mode of actions then synergy can be expected, and therefore, better eradication efficiency [19–21].

Biofilms are complex formations created by bacteria to improve their chances of survival. More specifically, biofilms can be 10 to 1000 times more tolerant than the planktonic phenotype [22]. The organization and composition existing in a biofilm makes the treatment of biofilm infections challenging since bacteria in biofilms can employ specific mechanisms to tolerate bactericidal treatments. The origin of biofilm tolerance is mostly caused by low metabolic activity of the bacteria within the biofilm, but it also has a genetic basis [23, 24]. Furthermore, the physical barrier of the biofilm matrix, limits the diffusion of molecules [25] into the biofilm, and reduces antimicrobial penetration.

Common sites of biofilm infections in the human body are the oral cavity, e.g. caries is the most frequent disease affecting human health [26], burdening billions of individuals with pain, limited masticatory functions and impaired aesthetics. In particular, the deep carious lesions [27], as well as its sequelae, the infected root canal associated with biofilm infection, represent targets for improved antimicrobial strategies and represent unsolved demanding challenges within the dental community [28]. Also within, the urinary system, the lungs of cystic fibrosis patients and chronic wounds biofilm infections are common. Additionally, when medical devices, like catheters, endoscopes, tissue fillers, implants, iatrogenic placed endodontic root filling materials, etc. are inserted into the body, the risk for chronic biofilm infection increases [29].

There is urgent need to enable elimination of chronic biofilm infections without utilizing excessive amounts of antibiotics. The present study could assist in developing light assisted combinatory treatments, consisting of irradiation in combination with antibiotics. The scope is to achieve total biofilm eradication and reduce the amount of antibiotics needed for treating infections.

In the present work we aimed to demonstrate the wavelength dependent survival of 24 hours (h) grown *P. aeruginosa* biofilms in the range 249 nm to 338 nm with an approximate step of 5 nm. The photon rate was 0.0036 mol/m^2^; corresponding to a radiant exposure of 1.700-1.260 J/m^2^. We report remarkable eradication (eradication log higher than 8) for the 24h old biofilms after irradiation in the 292-306 nm range for 0.0495 mol/m^2^ photon rate, corresponding to a radiant exposure 17.500-21.100 J/m^2^. A UVC treatment at that exposure level could have negative implications to the healthy tissue infected by the biofilm; therefore, the wavelengths tested at this higher level of radiant exposure were restricted only to the UVB and UVA region.

To demonstrate the antimicrobial effect of the irradiation method, we compared the irradiation strategy with two types of antibiotics that are well recognized for their usage against *P. aeruginosa* biofilm infections, namely tobramycin [30] and colistin [31] both as monotherapy. The hypothesis was no difference of the eradication efficacy of the three different treatments: UVB or antibiotic monotherapies, on *in vitro P. aeruginosa* biofilms either grown for 24h or 48h.

Light assisted antibiotic treatments for biofilm infections could be a method to improve the therapy of biofilm infections in the future, since light has been shown to have antibacterial action at several wavelengths [7]: UVC, UVB, UVA, blue, infrared. Here, we demonstrate the light assisted antibiotic principle in specific, by applying irradiation with a UV LED exhibiting central wavelength at 296 nm and by subsequently administrating topically tobramycin, to combat bacteria of *P. aeruginosa* biofilms grown for 24h (immature) or 48h (mature). The biofilm eradication of the light assisted tobramycin is compared to the effect achieved by 10fold more concentrated tobramycin, as well as combinatory antibiotic treatments, consisting of tobramycin and colistin at two concentration levels. Tobramycin and colistin are known to be used in combination due to the increased effectiveness of the combined treatment compared to monotherapy [18]. The hypothesis was no difference of the eradication efficacy of the four different treatments: light assisted tobramycin or 10-fold more concentrated tobramycin monotherapy and combinatory antibiotics at two different concentration levels, on in vitro *P. aeruginosa* biofilms either grown for 24h or 48h.

The proposed method is relevant for combating biofilms and could assist in developing new combinatory therapies consisting of light application and usage of antibiotics to improve treatment of chronic biofilm infections in complex ecosystems e.g. the dental root canal system [32].

## Results

### Wavelength dependent survival of biofilms

The survival of the biofilms after each treatment, was calculated according to Eq. 1, where N_treated_ is the number

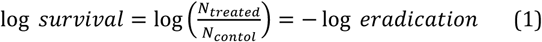

of colony forming units (CFUs) after a treatment is applied to the biofilm, and N_control_ is the number of CFUs of non-treated samples. The wavelength dependent survival of the *P. aeruginosa* biofilms grown for 24h is presented in Fig. 1. All treatments were repeated on different biological replicates. The statistical dispersion observed among technical replicates (orange error bars) was much smaller than the (statistical) dispersion observed among biological replicates. Therefore, the statistical dispersion reported in the following, measured as standard deviation, is of biological origin.

**Figure 1.**
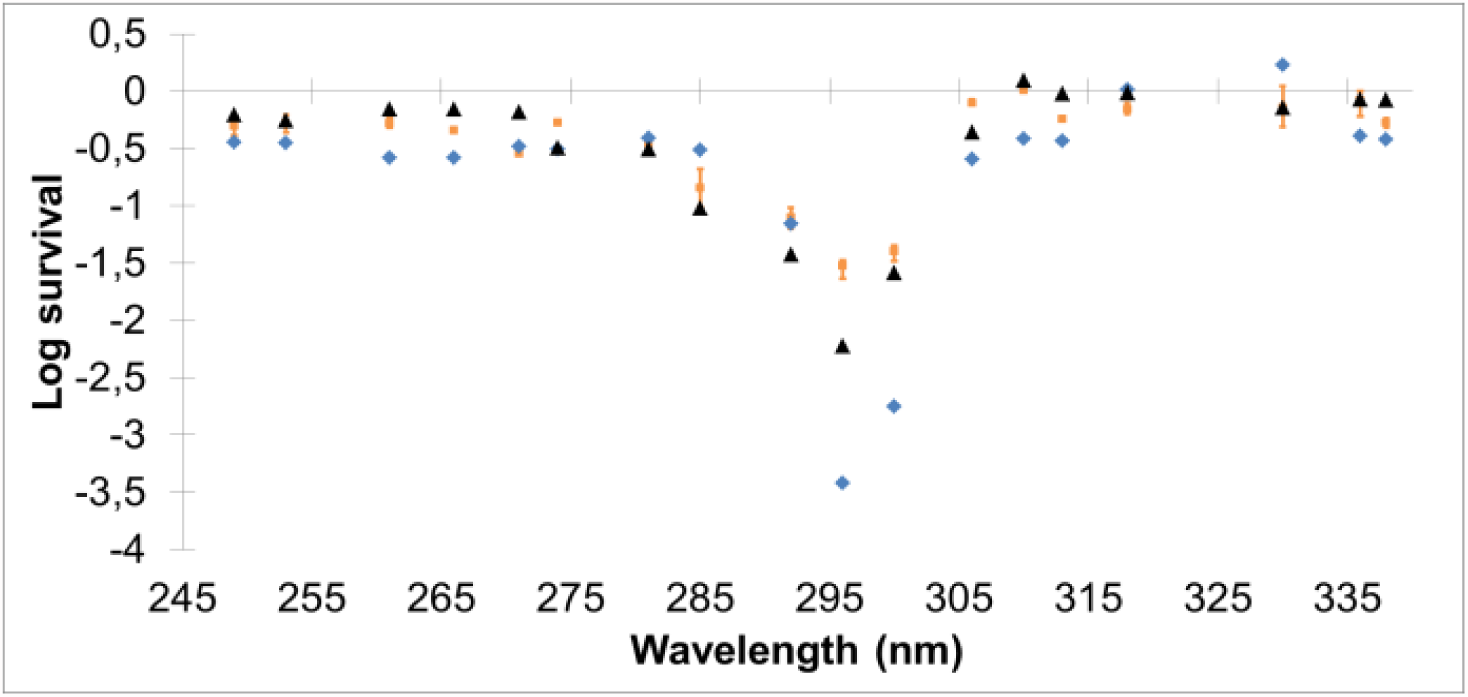
Wavelength dependent survival of *P. aeruginosa* biofilm grown for 24h. The photon rate delivered on the biofilms by the UV LEDs was 0.0036 mol/m^2^, corresponding to a radiant exposure 1.260 J/m^2^ to 1.700 J/m^2^, respectively, for the wavelengths 338 nm to 249 nm. Biological replicates were generated for all treatments as indicated by the different colors, orange, black and blue. For one of the biological replicates (orange) also technical replicates were generated; the error bars represent propagation of uncertainty for the function of log survival, coming from the different technical replicates (both for the treated and control samples).

UVA irradiations were performed with LEDs having central wavelengths from 318 nm to 338 nm, UVB irradiations from 281 nm to 313 nm, covering the whole UVB spectral range, and UVC irradiations from 249 nm to 274 nm. It was observed that the estimated CFUs of the UVA treated samples, independently of wavelength applied, were similar to non-treated samples (control). The log survival was 0.13±0.17. Independently of wavelength, UVC treated samples exhibited a log eradication 0.36±0.15 (Eq. 1). UVB irradiated samples to the contrary exhibited strong dependence of; wavelength and eradication ability and less CFUs were observed, especially for the range 292 to 300 nm. The treatment with the; diode having central wavelength at 296 nm exhibited the strongest eradication potential on *P. aeruginosa* 24h grown biofilms with log eradication 2.39±0.78.

The eradication ability at a radiant UVB exposure, equivalent to 12 h of summer sunlight in Northern Europe, was remarkable (eradication log higher than 8) for the wavelength range 292-306 nm (Fig. 2). The observed result suggested that a UVB radiant exposure of that level and at this wavelength range could also enact eradication effects on mature biofilms that are known for their increased tolerance to antibacterial treatments.

**Figure 2.**
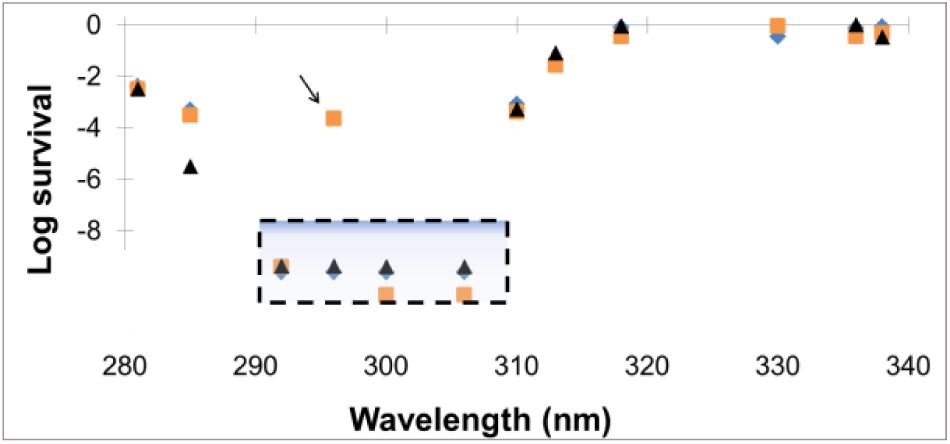
Survival of biofilm after UVB or UVA LED exposure, the level of exposure was equivalent to what can be received by sunlight. The photon rate delivered on the biofilms by the UVB and UVA LEDs was 0.0495 mol/m^2^ corresponding to a radiant exposure 17.500 J/m^2^ to 21.100 J/m^2^. The colors indicate different biological replicates. The biofilms were *P. aeruginosa* and grown for 24h. The peculiar observation indicated by the black arrow led us to perform several repetitions of this specific treatment; all repetitions resulted in log eradication higher than 8.

### UVB irradiation treatment versus topically administrated antibiotics

Three treatments were applied on *P. aeruginosa* biofilms grown for 24h or 48h, namely UVB irradiation (sunlight equivalent, 20.000 J/m^2^), or topical administration of antibiotics; either tobramycin or colistin at one hundred times the minimal inhibitory concentration (MIC). One-way analysis of variance (ANOVA) revealed that there was a significant difference in eradication of biofilms for the three treatments, both for 24h and 48h grown biofilms; the p. values were respectively 7.0 x **10 ^−4^** and 1.0 x **10 ^−5^**. The F. values with 8 degrees of freedom (Df1=2, Df2=6) were 30.9 and 127.7, respectively, for 24h and 48h grown biofilms.

The biofilm eradication achieved by the three different treatments is presented in Fig. 3. The eradication achieved by the UVB irradiation treatment was lower on mature biofilms (1.12±0.11 log eradication) than on immature biofilms (7.18±1.62 log eradication). However, the UVB treatment was more efficient in eradicating biofilms than the antibiotics. The colistin treatment resulted in negligible eradication, independently of the biofilm growth. The tobramycin treatment enacted a moderate eradication on immature biofilms (1.42±0.09 log eradication), but only negligible eradication on mature biofilms.

**Figure 3.**
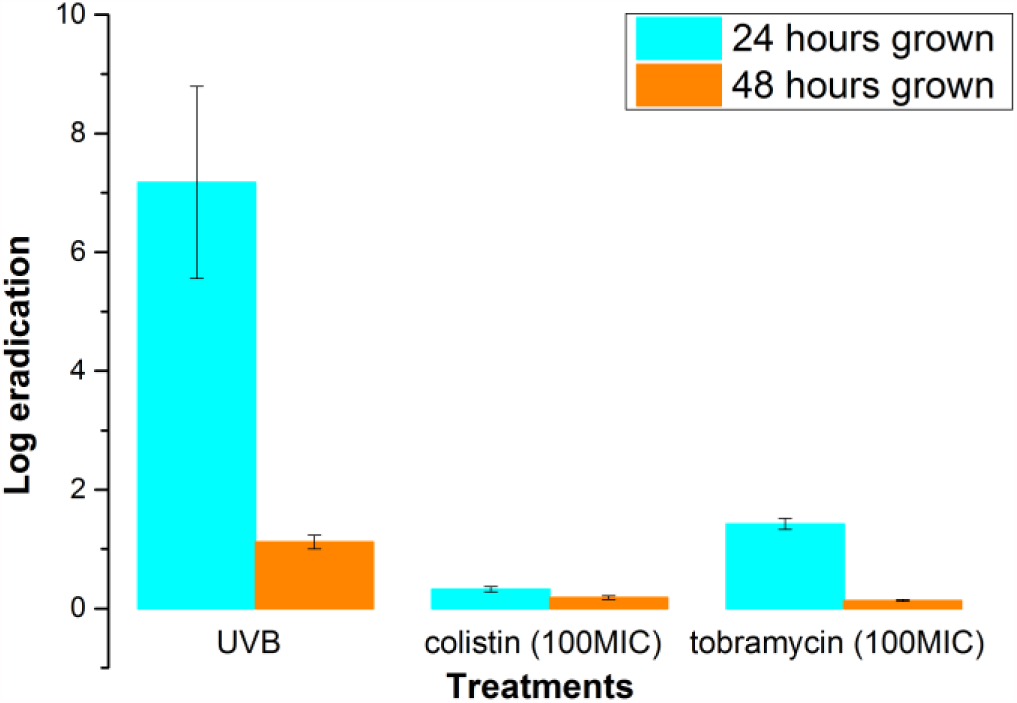
Eradication of biofilms after UVB irradiation treatment (sunlight equivalent), or topical administration of antibiotics; colistin or tobramycin at hundred times the MIC. The biofilms were either left to grow for 24h before treatment or for 48h. The error bars indicate the standard deviation as acquired by three biological replicates.

### Light assisted antibiotics

The biofilm eradication achieved by the suggested method of light assisted antibiotics is presented in Fig. 4. The eradication achieved by combinatory administration of antibiotics, and after administration of highly concentrated monotherapy is also shown in Fig. 4. The ANOVA analysis showed that there was a significant difference in the eradication achieved by the four treatments; both for 24h and 48h grown biofilms; the p. values were respectively 7.3 x **10 ^−8^** and 6.9 x **10 ^−6^**. The F. values with 11 degrees of freedom (Df1=3, Df2=8) were respectively 200.4 and 62.1 for 24h and 48h grown biofilms.

**Figure 4.**
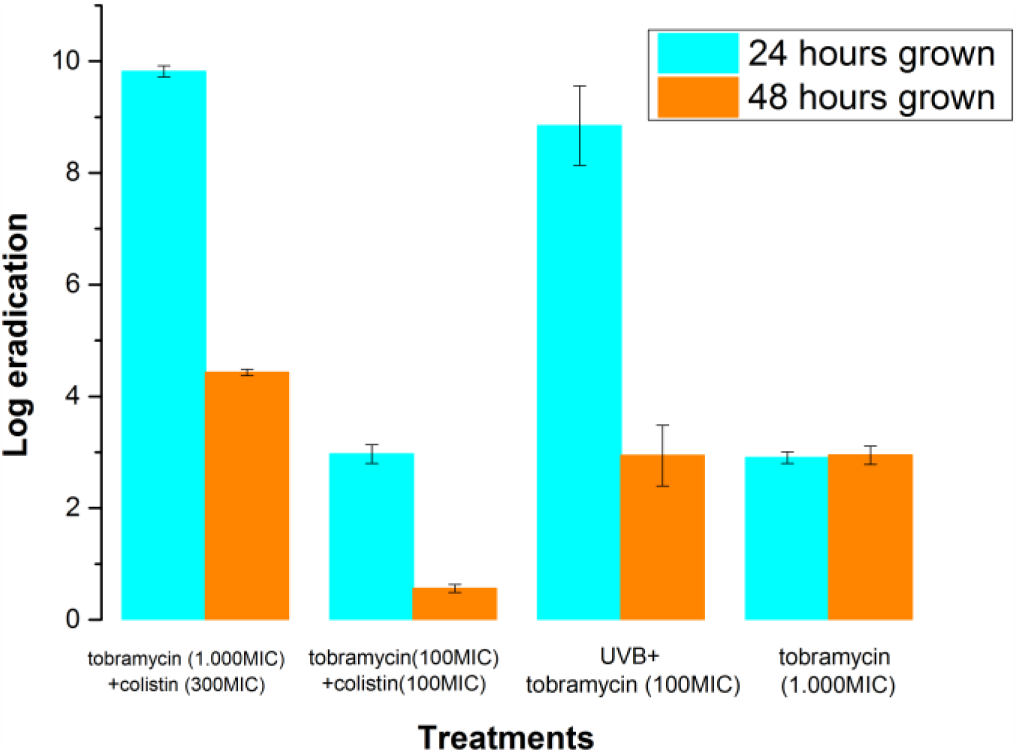
Eradication of biofilms achieved by light assisted tobramycin versus topical administration of 10-fold more concentrated tobramycin or combinatory antibiotics at two concentration levels. The biofilms were either grown for 24h or 48h before treatment. The error bars indicate the standard deviation as acquired by three biological replicates. “MIC” stands for minimal inhibitory concentration.

The light assisted tobramycin was even more effective than the combinatory antibiotics tobramycin and colistin (low-level concentration); and better than or equally as effective as high concentration of tobramycin depending on the age of the biofilm. Interestingly, the light assisted tobramycin treatment approached the eradication values achieved by topical administration of combinatory antibiotics at high concentrations; the light assisted tobramycin was 1 log less successful, both for combating 48h and 24h grown biofilms.

The log eradication values of the different treatments on 24h and 48h grown biofilms are presented in Table 1. Moreover, the 99% confidence limits, as determined by assuming a linear model for the treatments at each growth stage, are presented.

**Table 1.** Log eradication values after different treatments were applied on: 24h and 48h grown *P. Aeruginosa* biofilms. The 99% confidence limits; presented assume a linear model for the treatments, at each biofilm growth stage. “tobramycin+colistin high” stands for the combinatory antibiotic treatment, comprising of 1000 MIC tobramycin and ~300 MIC colistin. “tobramycin+colistin; low” stands for the combinatory treatment comprising of 100 MIC tobramycin and 100 MIC colistin.

| Growth of biofilm | Treatment | Log eradication | 0.5 % | 99.5 % |
| --- | --- | --- | --- | --- |
| 24 hours | Tobramycin+ colistin high | 9.82 | 8.57 | 11.06 |
|  | Tobramycin +colistin low | 2.97 | 1.72 | 4.21 |
|  | Light assisted tobramycin | 8.84 | 7.96 | 9.73 |
|  | Tobramycin monotherapy | 2.90 | 1.66 | 4.15 |
| 48 hours | Tobramycin+ colistin high | 4.43 | 3.47 | 5.39 |
|  | Tobramycin+ colistin low | 0.56 | 0.00 | 1.54 |
|  | Light assisted tobramycin | 2.94 | 2.26 | 3.62 |
|  | Tobramycin monotherapy | 2.95 | 1.98 | 3.91 |

## Discussion

*In vivo* and *in vitro* experiments have previously validated that the tolerance for biofilms are usually considerably higher (approx. 10-1000 times) than the planktonic bacterial cells [33]. Therefore, eradication of biofilms by conventional antibiotic administration can be challenging due to potential side effects or accumulated toxicity [34]. The demand for discovering alternative methods to eradicate biofilms, which may be involved in chronic infections, has been identified for a long time. Moreover, the need to develop treatments that would increase the vulnerability of biofilms to well-recognized therapeutic methods (e.g. antibiotics) has been acknowledged [35]. The established root canal infection associated with an apical periodontitis (i.e. inflammation surrounding the apical portion of the dental root) provides an area where light assisted antibiotics is applicable, as the prevailing combination of instrumentation and use of medicaments do not completely sterilize the root canal system [36].

Several bacterial species have previously been studied for their sensitivity towards UV irradiation, like *Escherichia coli* [37], and the optimal germicidal wavelength was found to lie within the UVC range with a maximum around 270 nm. However, the germicidal efficiency was studied for bacteria in the planktonic state. Bacteria in the planktonic state exist as individuals, while biofilms are aggregated bacteria. The peak of absorption of bacterial genetic material is located a few nanometers lower around 260-265 nm [38], and this supported the hypothesis that direct UVC absorption by bacterial genetic material, inhibits normal replication, and results in bacterial eradication [39–42].

In the present study it was shown that the eradication efficiency of UV irradiation on 24h grown *P. aeruginosa* biofilms is wavelength dependent, and that the optimum region is located in the UVB range around 296 nm (292-300 nm). The maximum optical thickness for the 24 h grown *P. aeruginosa* biofilms treated in the present work was 75±17 μm and for the 48 h grown 104±12 μm. The maximum physical thickness was respectively 100±23 μm and 138±16 μm for the 24 and 48 h grown biofilms.

In the biofilm state, the smaller penetration achieved by shorter wavelengths is expected to reduce the eradication efficacy of UVC [6]. In the UVA region and longer wavelengths bacterial eradication is dictated only by indirect pathways like generation of reactive oxygen species, and therefore, the eradication efficiency is much lower [43]. UVB is located spectrally between the UVC and UVA regions; and involves elements from both indirect and direct bacterial impairment [44, 45]. Studies on UVB lethality and mutagenesis of bacterial suspensions have shown that lethality occurs at a few nanometers longer wavelengths than mutagenesis [46]. Photons with UVB wavelengths in the 292-307 nm interval are expected to bring enough energy to break bonds like C-H and N-H [47], essential for the tertiary structure of proteins and DNA [48]. In the human skin, free radical generation exhibits high efficiency for wavelengths around 303 nm (UVB range) and 355 nm (UVA range) [49]. Recently, a product with strong absorbance at 297nm was reported by Puri *et al.* [50] as present in *Methylobacter tundripaludum* supernatants in a quorum sensing dependent manner. The collected product was reported to have no distinguishable growth inhibitory activity against E.coli MG 1655 or *Bacillus subtilis PY79*, however, a possible growth inhibitory action of the product was not excluded for other bacterial species.

The route of antibiotic administration in the present study was that antibiotics were added directly to the biofilm. Therefore, the biofilm should be directly accessible to the antibiotic administration. It was demonstrated that when tobramycin at a concentration, which only imposed negligible eradication effect on 48h grown biofilms, was administrated after UVB irradiation; it caused much larger eradication efficacy (2.94±0.54 log eradication) and reached the same eradication values as 10-fold more concentrated tobramycin. The eradication effect from UVB alone on 48h grown biofilms was significantly lower (1.12±0.11 log eradication). This indicates a synergetic effect of light and antibiotics of which the exact mechanism remains to be understood and optimized according to the taxonomic diversity of the biofilm to be eradicated.

## Conclusion

In conclusion, we have tested the efficiency of UV irradiation treatments to eradicate *P. aeruginosa* biofilms grown for 24h in the wavelength range 249 nm to 338 nm with an approximate step of 5 nm. It was shown that the log survival of the biofilm was remarkably reduced for the wavelength range 292-306 nm, and the optimum was located at 296 nm. Moreover, we demonstrated that the UVB irradiation was more efficient than topical administration of antibiotics (colistin or tobramycin at 100 MIC) for eradicating biofilms grown for 24h or 48h.

A novel method was introduced, light assisted antibiotics, for eradicating mature biofilms and successfully reducing the amount of antibiotics used for disinfection. A specific light assisted antibiotic example was presented; namely, irradiation with a UV LED exhibiting central wavelength at 296 nm combined with topical administration of tobramycin at 100 MIC. This treatment reduced the bacterial load on 48h grown biofilms by 3 logs, equivalent effect as that achieved by administrating 10-fold more concentrated tobramycin (1.000 MIC). The eradication achieved by the treatment was observed to be more effective than combinatory antibiotic treatment, 100 MIC of colistin plus 100 MIC of tobramycin. The light assisted tobramycin at 100 MIC was 1 log less successful than the high concentration combinatory antibiotic treatment, 1.000 MIC tobramycin plus 300 MIC of colistin. The present study can assist in developing new combinatory treatments consisting of light and usage of antibiotics to improve treatments of chronic biofilm infections within chronic wounds or within the infected root canal system treating infections in the jaw.

## Methods

### UV Irradiations

The UV LEDs that were used to perform the irradiation treatments were purchased from Sensor Electronic Technology, Inc (SETi, Columbia, SC, USA). The spectral irradiance of the diodes used to determine the optimal biofilm eradication wavelength, is depicted in Fig. 5. The spectral irradiance was measured by an External Optical probe (EOP-146, Instrument Systems GmbH, Munich, Germany) and a monochromator. The spectrometer used was a SPECTRO 320 (D) Release 5 (Instrument Systems GmbH). The exact protocol for measuring the spectral irradiance can be found in Barnkob et al [51]. The irradiance delivered on the biofilms was measured with a portable radiometer (NIST Certified UV Radiometer) at a distance (1.5±0.1cm). The distance between the biofilms and the UV LEDs was 1.5±0.2cm for all exposures; the error originates from the agar height, on which the filter carrying the biofilm was placed. The biofilms were kept in a UV free environment, when not treated.

**Figure 5.**
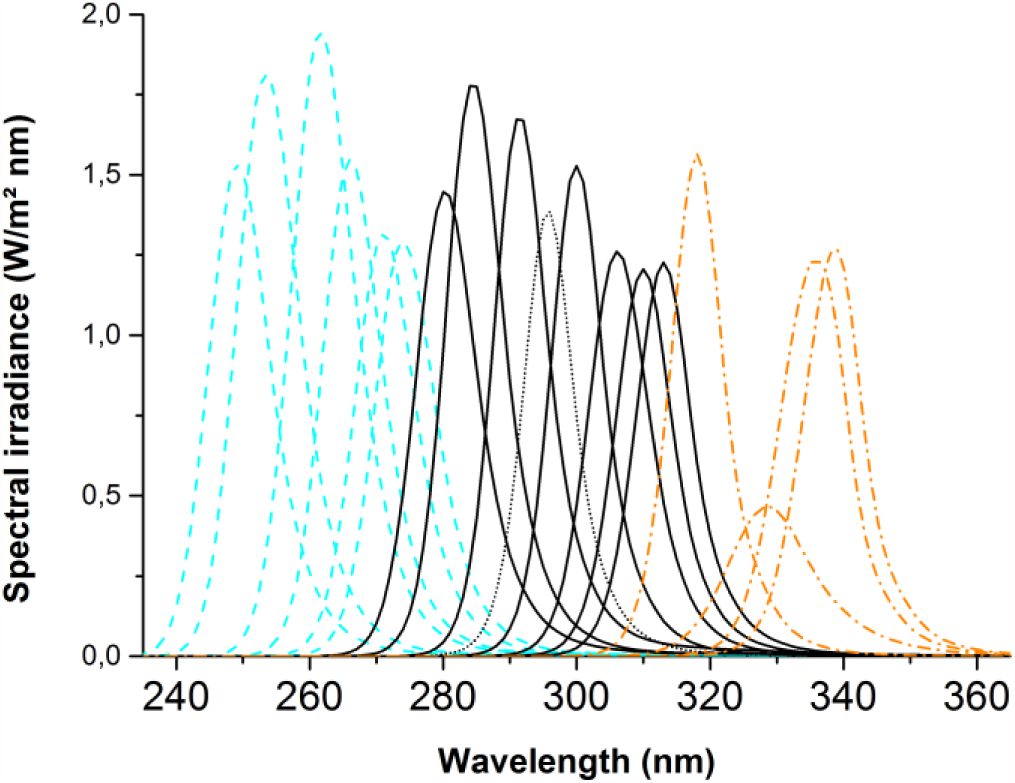
Spectral irradiance of UV LEDs used for determination of the optimal biofilm eradication wavelength. The UVC diodes are indicated with cyan color and dash line. The UVB diodes are indicated by black line, and black short dash for the optimal diode, central wavelength at 296nm. UVA diodes are indicated by orange dash dot line.

### Antibiotic treatments

The antibiotics used in this study were tobramycin (Eurocept International, Netherlands) and colistin sulfate salt (Sigma-Aldrich, USA). The MIC concentration for colistin was determined as 0.8 μg/mL and for tobramycin 1 μg/mL. The antibiotic treatments delivered were: 100 μg/mL tobramycin (100 MIC); 80 μg/mL colistin sulfate salt (100 MIC); tobramycin and colistin high (tobramycin+colistin high): 1000 μg/mL tobramycin (1.000 MIC) and 250 μg/mL colistin sulfate salt (~300 MIC); tobramycin and colistin low (tobramycin+colistin low): 100 μg/mL tobramycin and 80 μg/mL colistin sulfate salt; 1.000 μg/mL tobramycin (1.000 MIC). For the light assisted antibiotic treatment; 100 μg/mL tobramycin were delivered after UVB was applied (sunlight equivalent, 20.000J/m^2^). The antibiotics were added into the ABTG agar plates and biofilms were treated by moving the biofilm growing on nitrocellulose filters to the antibiotic containing media plates.

### Biofilm preparation

The bacterial strain used at the experiments was *P. aeruginosa* PAO1 [52]. A detailed description about the biofilm preparation can be found in Argyraki *et al.* [6]. The micropore assay methodology used to form the biofilms can be found in Bjarnsholt *et al.* [53]. Cellulose nitrate membrane filters with pore size 0.2μm and diameter 25 mm were purchased from Whatman GmbH (Germany). For biofilms grown for 48h, the membrane filters containing the biofilm were transferred every 24h onto fresh media to enable growth.

### CFU determination

The method for CFU determination can be found in Argyraki et al [6]. The serial dilutions in the cases of UVB irradiation treatment versus topically administrated antibiotics and light assisted antibiotics were performed from 10^0^ (no dilution) to 10^−7^ and the spotted volume was 10 μL; resulting in a detection limit of 100 bacteria per ml.

### Statistics

All treatments were performed on three different biological replicates (n=3) as a standard for testing reproducibility. The statistical dispersion is measured as standard deviation, reported in errors, and is of biological origin if not stated otherwise in the text. ANOVAs were two tailed and the software used for performing the statistical analysis was R [54]. In particular the function lm [55, 56] was used for determining the 99% confidence limits.

## Acknowledgements

The authors thank Anne Nielsen for her help in the laboratory and Sabrina Gericke for the blind counting of the CFUs. For financial support we would like to thank Region Zealand in Denmark (“the photonics green lab DOLL”, grant number: 2075008)

## Author contributions

All authors discussed the design of the study, the results, and contributed to the preparation of the manuscript. A.A. analyzed the data, performed irradiation treatments and wrote the majority of the manuscript. P.M.P. conceptualized the irradiation treatments.

M.M. grew the biofilms, analyzed data and performed the serial dilutions; for the wavelength dependence related data. C.S. grew the biofilms and performed the serial dilutions for the comparisons to antibiotics and light assisted treatments; and performed the antibiotic treatments. K. N. K. performed the biofilm thickness measurements. L.B. and M.M. designed the dental aspects of the study. T.B. designed the microbiological aspect of the study, analyzed microbiological data, as well as assisted in writing the microbiology part of the manuscript.

## Materials & Correspondence

Correspondence and material requests should be addressed to P.M.P and A.A.

